# Time-Resolved Phenotypic and Transcriptomic Responses of Primary Canine Dermal Fibroblasts to Prolonged Hypothermic Stress

**DOI:** 10.64898/2026.08.14.744362

**Authors:** Yuhan Wang, Ellie Shen, Alicia Huang, Elliot Lu, Yaxuan Liu, Jenny Huang, Bing Yu, Qingqing Dai

## Abstract

Prolonged low-temperature exposure may extend the preservation window of mammalian cells but can also disrupt cellular homeostasis and ultimately compromise cell viability. This study investigated the time-dependent phenotypic and transcriptomic responses of primary canine dermal fibroblasts to sustained hypothermic stress. Passage-three fibroblasts were continuously maintained at 15 °C for up to 15 days, with samples collected on Days 0, 3, 6, 9, 12, and 15. Cellular morphology, metabolic activity and viability, and apoptosis were evaluated using bright-field microscopy, Cell Counting Kit-8 assays, and Annexin V-FITC/propidium iodide flow cytometry, respectively. RNA sequencing was performed to characterize dynamic transcriptional changes throughout the exposure period. Early low-temperature exposure was associated with relatively preserved cellular morphology and viability, suggesting a transient adaptive response. With increasing exposure duration, fibroblasts exhibited progressive morphological deterioration, reduced metabolic activity, loss of adhesion, and increased apoptosis. Time-series transcriptomic analysis further revealed temporally coordinated and stage-dependent gene-expression programs associated with metabolic regulation, cellular stress responses, structural homeostasis, and cell survival. Integration of phenotypic and transcriptomic data demonstrated that the response of primary canine dermal fibroblasts to 15 °C was dynamic rather than linear, progressing from early adaptation to cumulative dysfunction during prolonged exposure. These findings provide a framework for defining the low-temperature tolerance of primary canine dermal fibroblasts and may inform the optimization of protocols for their short- to medium-term preservation and transportation.

## 1. Introduction

Dermal fibroblasts represent the principal resident mesenchymal cell population of the dermis and are indispensable for maintaining tissue homeostasis, synthesizing and remodeling the extracellular matrix, and orchestrating cutaneous wound repair[1,2]. Upon tissue injury, these cells are activated by a complex interplay of biochemical cues and mechanical stimuli, which triggers a coordinated response involving proliferation, directed migration into the wound bed, de novo synthesis of provisional matrix components, and ultimate differentiation into contractile myofibroblasts[3,4]. Dysregulation at any stage of this highly orchestrated repair cascade can arrest the healing process, leading to chronic non-healing wounds, or conversely, drive excessive ECM deposition and pathological fibrosis[5]. The dog has emerged as an increasingly relevant large-animal model in cutaneous research, as canine skin shares key physiological and pathological features with human skin that are challenging to replicate fully in conventional rodent models, which rely predominantly on wound contraction via the panniculus carnosus[6]. Moreover, naturally occurring diseases in companion dogs offer unique and valuable opportunities for comparative and translational research, serving as a critical bridge between induced small-animal disease models and human clinical trials[7]. Consequently, primary canine dermal fibroblasts serve as an important experimental platform for investigating fundamental skin physiology, mechanisms of wound repair, regenerative medicine strategies, and the development of standardized protocols for the ex vivo preservation of veterinary cell products[8,9].

Temperature is a fundamental environmental determinant of mammalian cell physiology[10]. Reducing culture temperature can suppress enzymatic reactions, energy consumption, cell-cycle progression, and overall metabolic activity, and has therefore been explored as a strategy for extending the storage and transportation window of cells and tissues[11]. Nevertheless, exposure to temperatures substantially below the physiological range can also disrupt cellular homeostasis[12]. Hypothermic stress has been associated with alterations in membrane fluidity, ion gradients, mitochondrial function, redox balance, protein folding, and cytoskeletal organization[13,14]. In particular, the accumulation of reactive oxygen species during hypothermia or subsequent metabolic imbalance may damage cellular macromolecules and activate apoptotic or necrotic pathways[15,16]. Thus, although moderate hypothermia may transiently reduce metabolic demand, its eventual biological consequences depend strongly on temperature, exposure duration, cell type, and the physiological state of the cells.

Cellular responses to low temperature are not necessarily linear[12]. Mammalian cells can initiate adaptive programs—including metabolic suppression, stress-response signaling, and remodeling of RNA and protein synthesis—during the early phase of cold exposure[17,18]. With prolonged stress, however, the capacity of these protective mechanisms may be exceeded, resulting in proliferative arrest, loss of adhesion, mitochondrial dysfunction, and irreversible cell death[19–22]. Primary cells may be especially susceptible because they retain donor-derived physiological characteristics and have a more limited proliferative capacity than immortalized cell lines. Although temperature-dependent growth regulation has been extensively investigated in established mammalian cell lines, relatively little is known about the long-term response of primary canine dermal fibroblasts to continuous culture at 15 °C. In particular, the temporal transition from early adaptation to progressive structural and functional injury has not been systematically characterized in this cell type.

In the present study, primary canine dermal fibroblasts were continuously cultured at 15 °C for up to 15 days to characterize their time-dependent phenotypic and transcriptomic responses to prolonged low-temperature stress. Samples were collected on Days 0, 3, 6, 9, 12, and 15 for bright-field morphological observation, CCK-8-based assessment of cellular metabolic activity and viability, Annexin V/propidium iodide flow-cytometric analysis of apoptosis, and RNA sequencing. By combining these complementary approaches, we sought to define the temporal boundary between short-term cellular adaptation and progressive injury and to identify both conserved and stage-specific transcriptional programs associated with sustained hypothermic stress. This study provides a cellular and molecular framework for understanding the low-temperature tolerance of primary canine dermal fibroblasts and may facilitate the development of improved protocols for their short- to medium-term preservation, transportation, and experimental use.

## 2. Results

### 2.1. Prolonged 15 °C Culture Induces Time-Dependent Morphological Damage, Proliferative Suppression and Apoptosis in Canine Dermal Fibroblasts

To systematically characterize the biological responses of cells subjected to sustained low-temperature stress at 15 °C, we performed continuous long-term culture for up to 15 days and collected specimens at Day 0, 3, 6, 9, 12 and 15 for bright-field morphological observation, CCK-8 cell viability assay, and Annexin V/PI double-staining flow cytometric apoptosis detection.

Bright-field microscopic imaging revealed gradual and aggravated morphological lesions in cells upon prolonged incubation at 15 °C (Figure 1A). Cells cultured at Day 0 displayed typical elongated spindle shape, intact cell membrane and tight adherent state, representing normal cellular morphology. No obvious structural damage was observed on Day 3 and Day 6. Starting from Day 9, partial cellular shrinkage, cytoplasmic vacuolization and increased cell refractivity emerged in a fraction of cells. More severe pathological changes were observed at Day 12, including obvious cell rounding, cytoplasmic fragmentation and appearance of suspended dead cell debris. At the endpoint of Day 15, the majority of adherent cells exhibited severe shrinkage, spherical transformation and massive detachment from the substrate, indicating irreversible cell injury triggered by long-term 15 °C culture stress.

**Figure 1.**
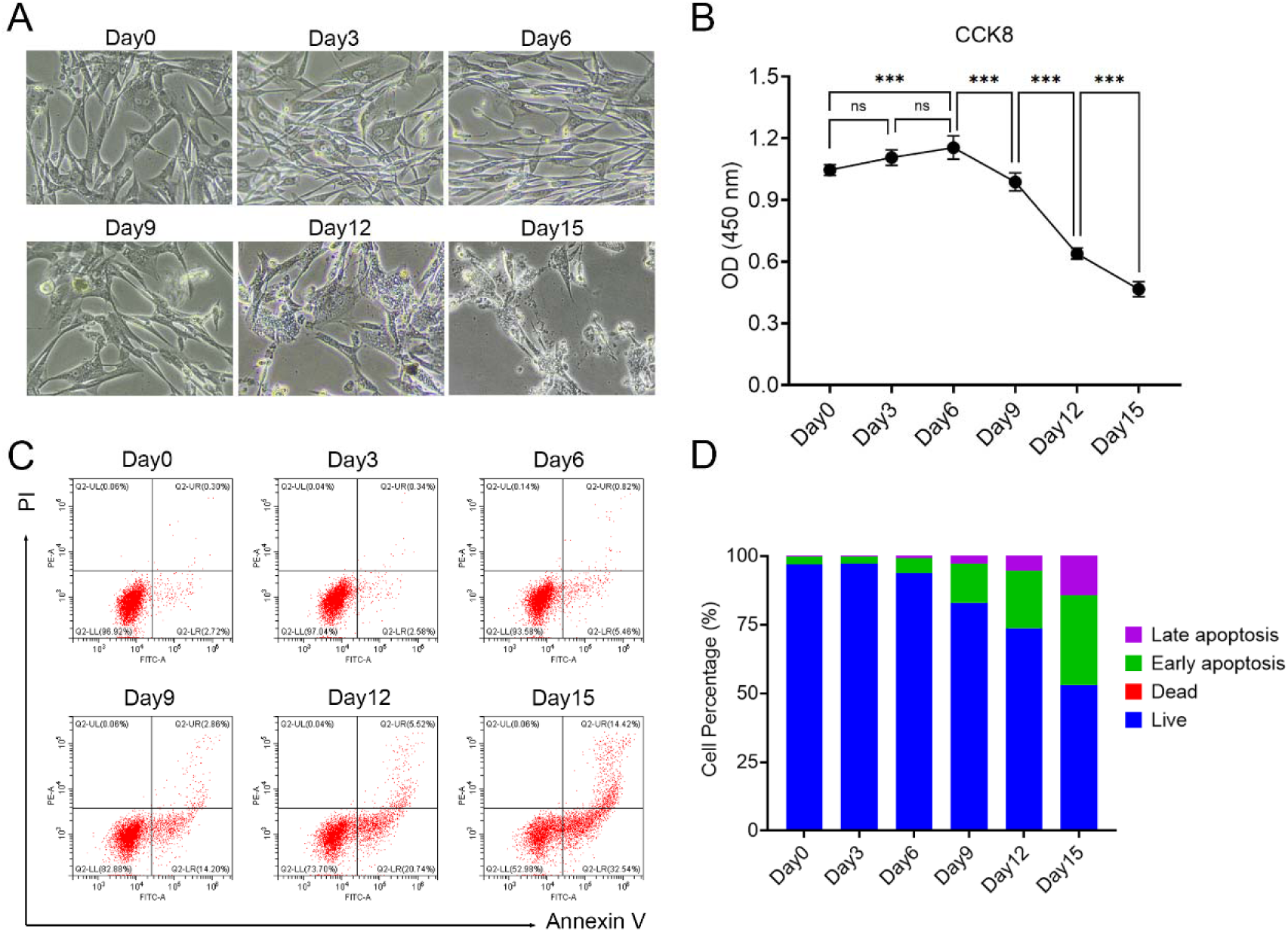
Morphological, proliferative and apoptotic changes of cells after continuous long-term culture at 15 °C for 0–15 days. (A) Bright-field microscopic images showing cell morphological alterations at Day 0, 3, 6, 9, 12 and 15 under 15 °C prolonged incubation. (B) Metabolic activity and viability were determined by CCK-8 assay, presented as OD values at 450 nm at different time points; error bars indicate mean ± SD. Statistical analysis was performed via one-way ANOVA followed by Dunnett’s test. ns, not significant; ***, P < 0.001. (C) Representative flow cytometry dot plots of Annexin V-FITC/PI double staining for apoptosis detection. Quadrant definition: Q2-LL, live cells; Q2-LR, early apoptotic cells; Q2-UR, late apoptotic cells; Q2-UL, necrotic/dead cells. The percentage of each cell subset is marked in corresponding quadrants. (D) Stacked bar graph for statistical quantification of live cells, early apoptotic cells, late apoptotic cells and dead cells based on flow cytometric results.

A CCK⍰8 assay was performed to quantify cell metabolic activity and viability across all time⍰point groups (Figure 1B). The absorbance value at 450 nm increased mildly from day 0 to day 6 and peaked at day 6, reflecting short term metabolic adaptation of cells upon early⍰phase low⍰temperature stress. Subsequently, optical density values dropped steadily between day 6 and day 15, with pronounced reductions observed on day 9 and day 12; cell viability bottomed out on day 15. This declining trend suggests that sustained incubation at 15 °C produces time⍰dependent cumulative suppression on cellular metabolism, which progressively impairs cell viability with prolonged cold exposure.

We further adopted Annexin V-FITC/PI double-labeled flow cytometry to quantify the proportions of viable cells, early apoptotic cells, late apoptotic cells and necrotic dead cells (Figure 1C, D). Representative dot plots and stacked bar statistical histogram showed that cells remained almost fully viable with minimal apoptotic populations from Day 0 to Day 6. As the culture duration extended to Day 9 and beyond, the percentage of live cells decreased in a time-dependent manner, accompanied by synchronous elevation of early apoptosis (Annexin V⁺PI⁻) and late apoptosis (Annexin V⁺PI⁺) cell fractions. At Day 15, viable cells only accounted for approximately 50% of the total cell population, while early and late apoptotic cells occupied substantial proportions. These flow cytometry data quantitatively verified that long-term incubation at 15 ° C triggered time-course progressive apoptotic cascade activation and viability loss in the tested cells. In summary, sustained 15 °C low-temperature long-term culture leads to progressive morphological destruction, proliferation arrest and dose-time-dependent apoptosis in adherent cells.

### 2.2. Time⍰resolved Transcriptomic Remodeling Under Sustained 15 °C Hypothermic Stress

To systematically dissect the dynamic transcriptomic landscape of cells under sustained 15 °C low-temperature stress across 0, 3, 6, 9, 12 and 15 days of continuous culture, we performed RNA-sequencing (RNA-seq) on samples from all six time points.

PCA based on all gene expression profiles was applied to evaluate the overall transcriptional similarity and divergence of samples at different culture time points (Figure 2A). The first principal component (PC1) explained 71.58% of total transcriptional variance, while the second principal component (PC2) accounted for 19.56% of variance. Clear time-dependent clustering and separation were observed: Day 0 samples clustered independently and were distinctly distant from all later time points. Samples from Day 3 and Day 6 aggregated into adjacent clusters, whereas Day 9, Day 12 and Day 15 were gradually dispersed along the PC1 axis with prolonged culture duration. This robust grouping pattern indicated that long-term low-temperature incubation triggered progressive, cumulative and time-ordered global transcriptomic remodeling, and the transcriptional profiles were gradually deviated from the baseline status of Day 0 over culture time.

**Figure 2.**
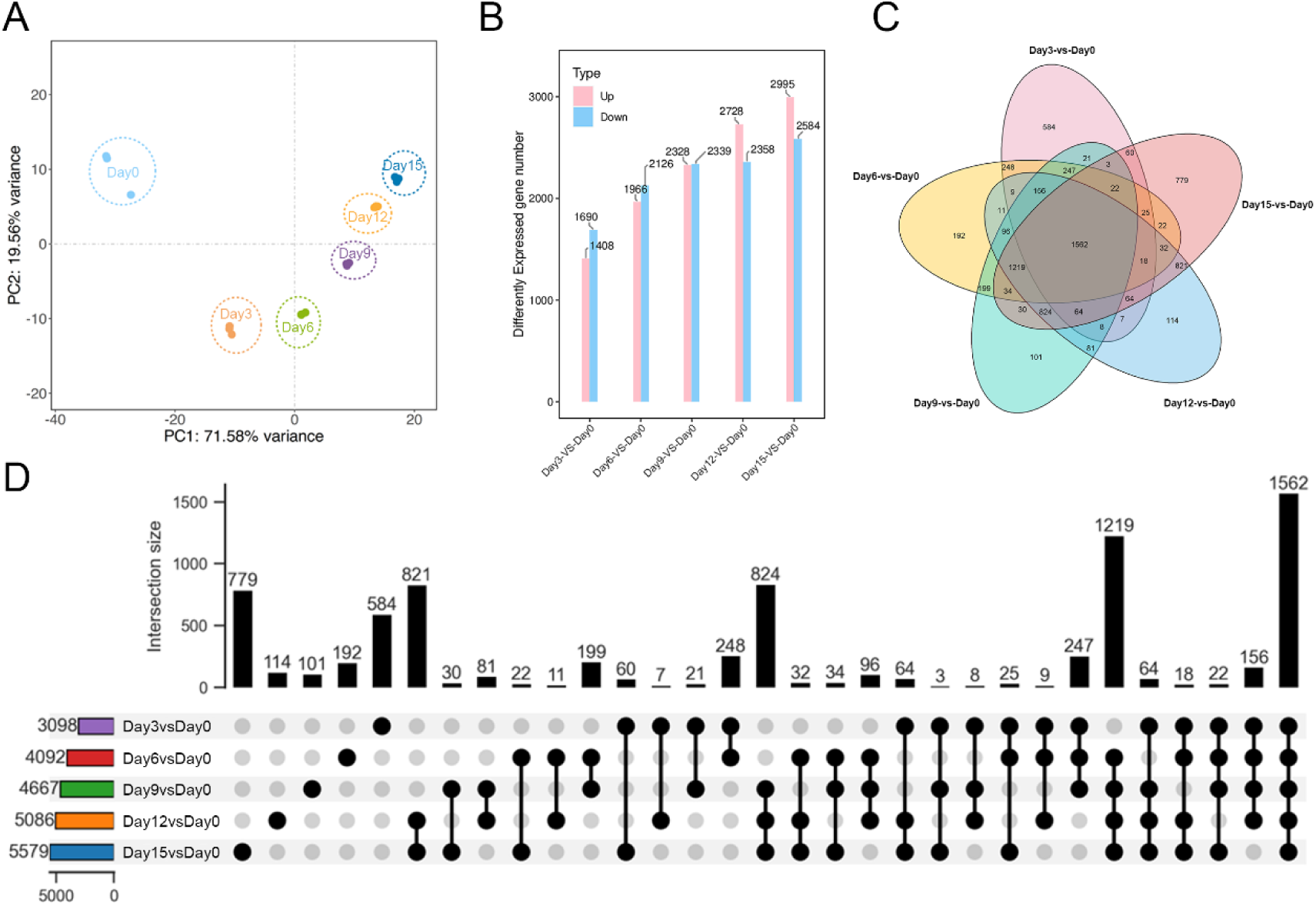
RNA-seq transcriptomic analysis of cells subjected to continuous 15 °C low-temperature culture for 0–15 days. (A) PCA plot showing global transcriptional clustering of samples collected at Day 0, 3, 6, 9, 12 and 15. PC1 and PC2 represent the top two principal components with explained variance labelled on axes. Each color circle corresponds to one culture time point. (B) Bar chart quantifying the number of upregulated (pink) and downregulated (blue) differentially expressed genes (DEGs) in each time point compared with Day 0 baseline. (C) Venn diagram illustrating the overlap and unique DEGs among five comparison groups (Day3 vs Day0, Day6 vs Day0, Day9 vs Day0, Day12 vs Day0, Day15 vs Day0). The number inside each region indicates the gene count of corresponding subset. (D) UpSet plot for quantitative statistical analysis of DEG intersections across all five pairwise comparisons. The vertical bars represent the number of intersecting genes, and the bottom dot matrix marks the comparison groups involved in each intersection.

We next identified differentially expressed genes (DEGs) for each time point by pairwise comparison against Day 0 baseline (Figure 2B). The number of both upregulated (pink) and downregulated (blue) DEGs exhibited a prominent time-dependent increasing trend with extended low-temperature treatment. Specifically, 1408 upregulated and 1690 downregulated genes were screened at Day3 vs Day0; the DEG quantity continuously elevated at Day6 (1966 up, 2126down), Day9 (2328 up, 2339 down), Day12 (2728 up, 2358 down), and reached the maximum at Day15, with 2995 upregulated genes and 2584 downregulated genes. The continuous expansion of DEG pool demonstrated that transcriptional perturbation was aggravated in a time-dependent manner under persistent 15 °C stress.

Venn diagram visually displayed the overlapping and unique DEGs across five comparison groups (Day3/6/9/12/15 vs Day0) (Figure 2C). A total of 1562 core shared DEGs existed in all five pairwise comparisons, representing the universal transcriptional signature persistently activated throughout the entire 15-day low-temperature culture process. UpSet plot was further employed to quantitatively and systematically quantify the intersection sizes of DEGs among all comparison sets (Figure 2D), which complemented and refined the Venn diagram results. Consistent with Venn analysis, the largest intersection set contained 1562 shared DEGs across all five groups. In addition, multiple intermediate overlapping subsets and time-point exclusive DEGs were clearly enumerated: the largest stage-specific DEGs were 779 unique genes in Day15 vs Day0, followed by 584 unique DEGs in Day3 vs Day0, 821 shared DEGs between Day12 and Day15, 824 shared DEGs between Day9, Day12 and Day15, etc. The UpSet statistics precisely quantified the magnitude of common responsive genes and time-restricted transcriptomic alterations, solidifying the conclusion that long-term low-temperature culture induced both universal core transcriptional programs and phase-specific gene expression changes during the progression of cellular stress response.

Collectively, transcriptomic data confirmed that prolonged 15 °C low-temperature culture drove gradual transcriptomic divergence, continuously expanded differential gene expression, and elicited both conserved core DEGs and time-stage-specific transcriptional signatures in stressed cells.

### 2.3. Temporal Clustering and Functional Annotation of DEGs in Response to Sustained 15 °C Hypothermia

To further dissect the dynamic expression patterns and biological functional implications of time-course DEGs triggered by long-term 15 °C cold culture, we performed STEM trend clustering for all DEGs, followed by KEGG pathway and GO biological process enrichment separately for significantly downregulated and upregulated gene sets.

STEM clustering classified all time-series DEGs into 8 distinct expression profiles with statistically significant enrichment (FDR < 0.05) (Figure 3A). Each module represented a unique temporal transcriptional trajectory from Day 0 to Day 15 during continuous low-temperature incubation. The top two largest clusters contained 2166 genes (continuously ascending expression trend) and 1486 genes (persistently descending expression trend), which constituted the two dominant transcriptional response modes under sustained cold stress. Other minor modules presented fluctuating expression patterns with peak or trough values at intermediate time points (Day 6, Day 9 or Day 12), implying staged and oscillatory gene regulation during chronic low-temperature stimulation. All clustered profiles exhibited extremely significant statistical reliability, confirming the robustness of these time-dependent gene expression trends.

**Figure 3.**
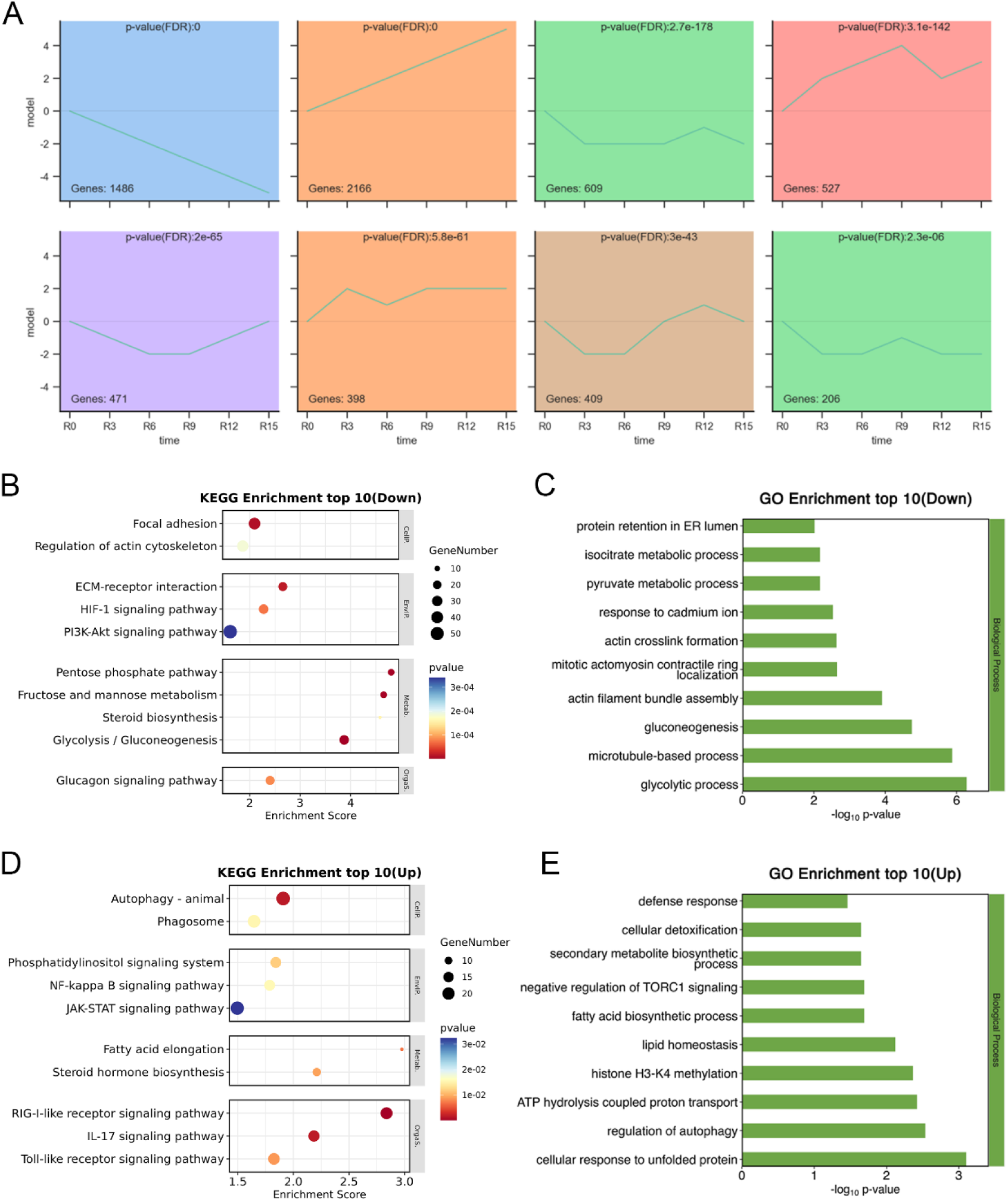
Time-series expression trend clustering and functional enrichment of DEGs upon long-term 15 °C low-temperature culture. (A) STEM clustering analysis showing eight statistically significant gene expression profiles across Day 0 to Day 15. The gene number within each module and adjusted FDR value are labelled for each trend cluster. X-axis: culture time points; Y-axis: normalized gene expression value. (B) Top 10 enriched KEGG pathways for significantly downregulated DEGs, presented as bubble plot sized by enriched gene number and colored by P-value. (C) Top 10 enriched GO biological process terms for downregulated DEGs, ranked by -log₁₀(P-value). (D) Top 10 enriched KEGG pathways for significantly upregulated DEGs in bubble plot format. (E) Top 10 enriched GO biological process terms for upregulated DEGs, ranked by -log₁₀(P-value).

Subsequently, KEGG pathway and GO enrichment analyses were performed on DEGs with continuously upregulated and downregulated temporal expression trends. KEGG enrichment revealed that downregulated genes were predominantly enriched in pathways closely associated with cell adhesion, cytoskeleton remodeling, energy metabolism and proliferation survival signaling, including focal adhesion, regulation of actin cytoskeleton, ECM-receptor interaction, HIF-1 signaling pathway and PI3K-Akt signaling pathway (Figure 3B). Multiple carbohydrate metabolic pathways were also markedly suppressed, such as pentose phosphate pathway, glycolysis/gluconeogenesis, fructose and mannose metabolism, which indicated impaired energy supply and glycolytic flux under long-term cold stress. Consistently, GO biological process enrichment for downregulated genes highlighted terms related to cytoskeleton assembly, contractile structure formation and central carbon metabolism: glycolytic process, gluconeogenesis, actin filament bundle assembly, mitotic actomyosin contractile ring localization, as well as endoplasmic reticulum protein retention (Figure 3C). These results collectively demonstrated that prolonged low-temperature culture broadly inhibited cell adhesion, cytoskeleton dynamics, PI3K/Akt survival signaling and core carbohydrate energy metabolism.

For significantly upregulated DEGs, top 10 KEGG and GO enrichment terms reflected stress-activated adaptive and defensive transcriptional programs (Figure 3D, 3E). KEGG pathway enrichment showed remarkable activation of autophagy-related cascades (autophagy-animal, phagosome), inflammatory and immune signaling axes (NF-kappa B, JAK-STAT, IL-17, Toll-like receptor, RIG-I-like receptor signaling pathways), together with lipid metabolism and steroid biosynthesis pathways (Figure 3D). GO biological process enrichment further validated the stress-responsive characteristics of upregulated genes, prominently enriched in defense response, cellular detoxification, cellular response to unfolded protein, negative regulation of TORC1 signaling, autophagy regulation, histone H3-K4 methylation and fatty acid/lipid homeostasis (Figure 3E). These functional annotations illustrated that chronic 15 °C low-temperature stress robustly activated autophagy, innate immune defense, protein homeostasis surveillance and lipid metabolic reprogramming as adaptive protective mechanisms in stressed cells.

In summary, time-series gene expression was grouped into multiple temporal trends by STEM analysis. Functional enrichment revealed that long-term cold stress repressed cell adhesion, cytoskeleton remodeling and glycolytic metabolism while strongly inducing autophagy, stress defense response, immune signaling and lipid metabolic adaptation at the transcriptome level.

### 2.4. GSEA Validation of Progressive Stress Pathway Activation and Metabolic Repression Under Sustained Hypothermia

To further verify the time-dependent pathway activity identified in the above KEGG enrichment results, we performed gene set enrichment analysis (GSEA) on the time-series transcriptome data and calculated normalized enrichment score (NES) for representative pathways at Day 3, 6, 9, 12 and 15 (Figure 4).

**Figure 4.**
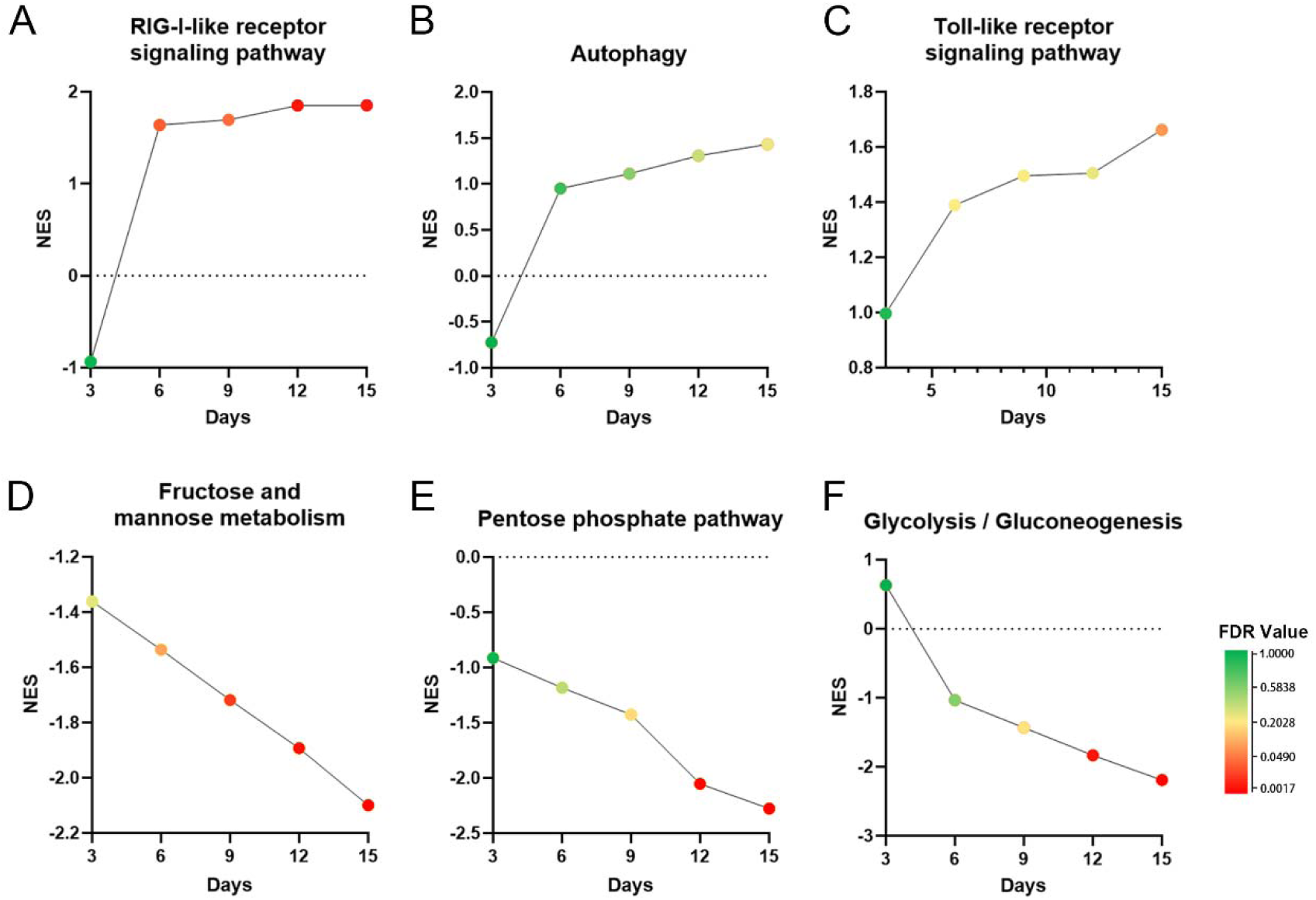
Dynamic changes in pathway activity evaluated by GSEA during prolonged low-temperature incubation. (A–C) Time-course normalized enrichment score (NES) of progressively activated pathways: (A) RIG-I-like receptor signaling pathway; (B) Autophagy; (C) Toll-like receptor signaling pathway. (D–F) Time-course NES of progressively repressed metabolic pathways: (D) Fructose and mannose metabolism; (E) Pentose phosphate pathway; (F) Glycolysis/Gluconeogenesis. The color of each data point corresponds to the FDR value, with green representing high FDR (low significance) and red representing low FDR (high significance). The dotted line at NES = 0 denotes no enrichment.

Consistent with the previous enrichment results, innate immune and autophagy pathways exhibited progressive activation as culture time extended (Figure 4A–C). The NES of the RIG-I-like receptor signaling pathway sharply increased from Day 3 and remained persistently elevated from Day 6 to Day 15 (Figure 4A). Similarly, the Autophagy pathway showed a continuous upward trend in NES, starting from a negative value at Day 3 and gradually rising throughout the 15-day culture (Figure 4B). The Toll-like receptor signaling pathway also displayed a steady increase in enrichment score, with its NES rising continuously and reaching the highest level at Day 15 (Figure 4C). The color gradient of data points further indicated that the FDR value gradually decreased (higher statistical significance) with prolonged incubation, confirming that these stress-responsive immune and autophagy cascades were progressively and significantly activated under chronic cold stress.

In contrast, carbohydrate metabolic pathways displayed robust and progressive transcriptional repression (Figure 4D–F). For Fructose and mannose metabolism, Pentose phosphate pathway, and Glycolysis/Gluconeogenesis, the NES values started relatively high at Day 3 and declined continuously over time, falling to strongly negative values by Day 15. Meanwhile, the FDR values gradually became smaller, demonstrating that the suppression of core carbon metabolism became increasingly significant as low-temperature stress persisted. These GSEA trajectories independently validated the findings from DEG enrichment analysis, demonstrating that long-term 15 °C culture chronically activates innate immune and autophagy programs while gradually inhibiting glycolysis and pentose phosphate flux at the transcriptional level.

### 2.5. Dynamic Expression Profiles and Validation of Core Hub Genes Under Sustained Hypothermic Stress

Based on the STEM trend clustering and functional enrichment results above, we further characterized the temporal expression patterns of two dominant gene modules: persistently downregulated and persistently upregulated DEGs across the 15-day low-temperature culture course, and verified the dynamic transcription of top continuously induced hub genes. Hierarchical clustering heatmaps visually presented the global expression trajectories of two major trend clusters (Figure 5A, 5B). Volcano plot of the Day15 vs Day0 comparison was used to locate the top 10 persistently upregulated core genes, including *PPP1R15A*, *ALAS1*, *AKAP8L*, *UBTD1*, *DPF2*, *SLCO2A1*, *DNAJC3*, *PLEKHM2*, *CCL5*, and *LASP1*, within the overall differential gene set (Figure 5C).

**Figure 5.**
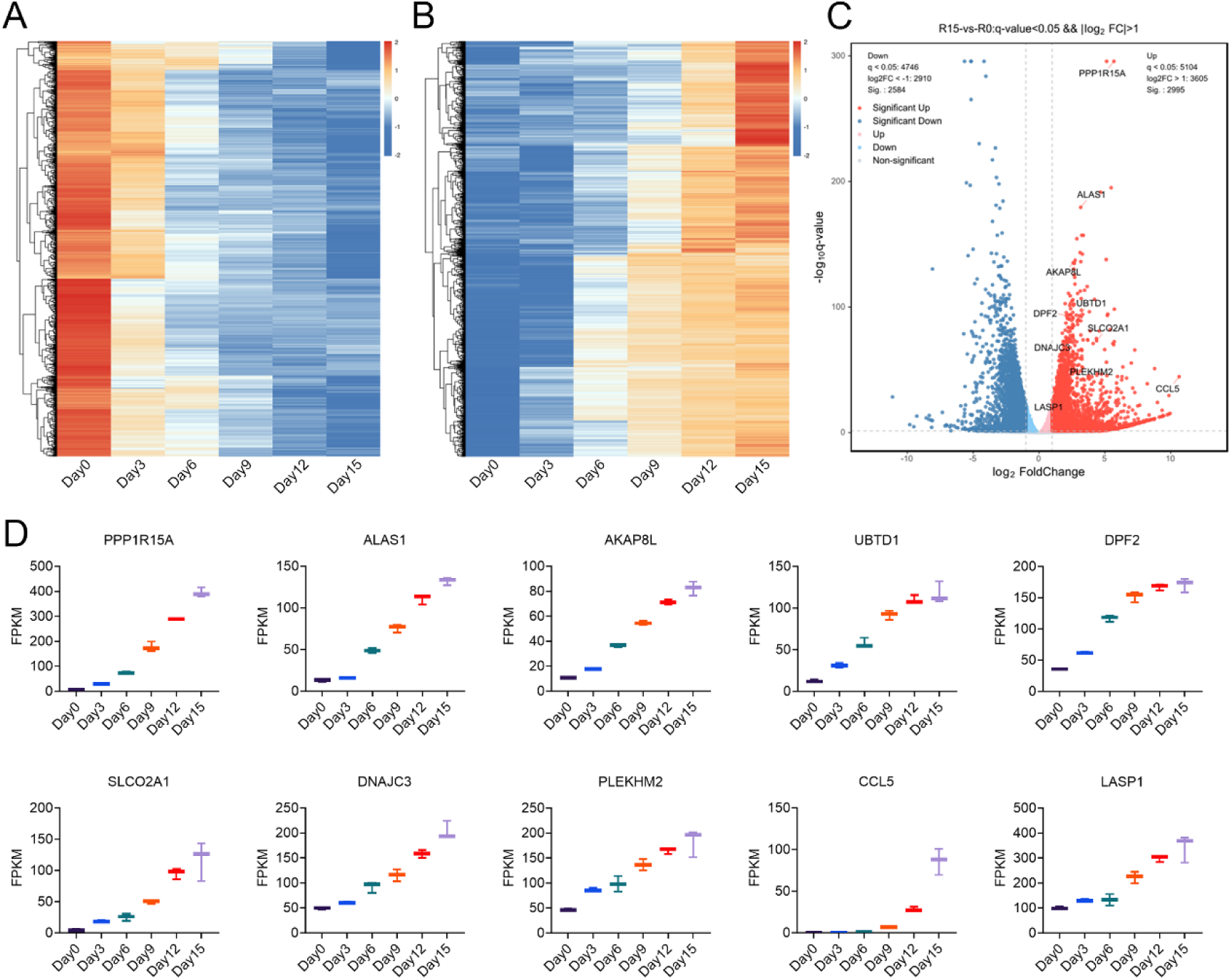
Temporal expression profiling of persistently regulated DEGs and validation of top upregulated hub genes. (A) Heatmap showing hierarchical clustering of genes with continuous downward expression trends from Day 0 to Day 15 under 15 °C culture. Rows represent individual genes, columns represent time points; color intensity indicates normalized expression level. (B) Heatmap of genes with continuous upward expression trends across the whole culture period. (C) Volcano plot of DEGs in Day15 versus Day0 comparison (threshold: q-value < 0.05, |log₂FC| > 1). Red dots denote significantly upregulated genes, blue dots denote significantly downregulated genes. The top 10 continuously upregulated hub genes are labelled. (D) Line/box plots showing dynamic FPKM expression values of the 10 core upregulated genes at six time points (Day 0, 3, 6, 9, 12, 15).

To intuitively validate their time-series expression trends, we plotted the FPKM abundance of these 10 hub genes at all six time points (Day 0, 3, 6, 9, 12, 15) (Figure 5D). All selected genes maintained extremely low basal expression at Day 0 and displayed a consistent stepwise upward tendency with extended low-temperature treatment, reaching the highest FPKM values at Day 15. Among them, PPP1R15A and LASP1 showed the most dramatic transcriptional induction.

In summary, long-term low-temperature stress drives two opposite large-scale transcriptional modules: a steadily declining gene set related to basic metabolism and cell structure, and a continuously rising stress-responsive gene cohort. Ten representative persistently upregulated hub genes were screened and verified by volcano plot and time-series FPKM quantification, serving as key candidate effectors mediating cellular adaptation to chronic cold challenge.

## 3. Discussion

The present study provides a time-resolved characterization of the phenotypic and transcriptomic responses of primary canine dermal fibroblasts exposed continuously to 15 °C for up to 15 days. By integrating morphological examination, CCK-8 metabolic activity analysis, Annexin V/propidium iodide flow cytometry, and RNA sequencing at six consecutive time points, the study reveals a biphasic response to sustained moderate hypothermia. During the first 3–6 days, the cells largely maintained their adherent spindle-shaped morphology, showed minimal apoptosis, and exhibited a modest increase in CCK-8 metabolic activity. In contrast, exposure for 9 days or longer was accompanied by progressive cell shrinkage, vacuolization, rounding, fragmentation, and detachment, together with reduced metabolic activity and increased apoptotic cell fractions. These phenotypic changes were paralleled by a pronounced time-dependent divergence of the transcriptome from the Day 0 state and an overall expansion of the differentially expressed gene pool. Collectively, these findings indicate that primary canine dermal fibroblasts can tolerate 15 °C for a limited period through an early adaptive response, but prolonged exposure eventually exceeds their homeostatic capacity and leads to cumulative, potentially irreversible cellular injury.

The preservation of cellular morphology and metabolic activity during the first 6 days suggests that moderate hypothermia does not cause immediate catastrophic damage in primary canine dermal fibroblasts. Lowering temperature generally decreases enzyme kinetics, membrane transport, protein synthesis, ATP turnover, and cell-cycle progression, thereby reducing the metabolic requirements of mammalian cells[23–25]. Such metabolic depression can temporarily preserve cellular resources and delay the accumulation of metabolic waste[26]. Mammalian cells may also activate cold-responsive mechanisms involving RNA-binding proteins, translational regulation, antioxidant defense, and remodeling of energy metabolism[26,27]. The slight increase in CCK-8 metabolic activity from Day 0 to Day 6 may therefore represent temporary maintenance or enhancement of cellular reducing activity during early adaptation. However, this result should not be interpreted unequivocally as increased cell proliferation. CCK-8 is based on the reduction of a water-soluble tetrazolium salt by cellular dehydrogenases and consequently reflects the combined effects of viable cell number and intracellular metabolic activity[28]. Direct proliferation measurements, such as cell counting, EdU incorporation, Ki-67 staining, or cell-cycle analysis, would be required to determine whether fibroblasts truly proliferated at 15 °C. Because mammalian cell-cycle progression is generally slowed at reduced temperatures, the early increase in absorbance may instead reflect enhanced metabolic activity per surviving cell, differences in attachment, or a transient compensatory response.

The transition observed from Day 9 onward is consistent with the cumulative nature of hypothermic injury. Low temperature initially suppresses metabolism but can simultaneously disturb membrane fluidity, ion homeostasis, cytoskeletal organization, mitochondrial respiration, and protein-folding processes[29,30]. Prolonged hypothermia can impair ATP production and the activity of energy-dependent ion pumps, resulting in ionic imbalance, cellular swelling or shrinkage, and loss of membrane integrity[31–33]. Mitochondrial dysfunction and disturbed electron transport may also promote the generation of reactive oxygen species, whereas antioxidant and repair systems can become less effective under sustained metabolic stress[34,35]. The resulting oxidative imbalance can damage lipids, proteins, and nucleic acids and initiate mitochondria-dependent apoptotic signaling[36]. The emergence of cytoplasmic vacuolization and enhanced refractivity at Day 9 may therefore represent an intermediate injury phase in which stress adaptation is no longer sufficient to maintain normal cellular architecture. The more severe rounding, fragmentation, and detachment observed on Days 12 and 15 indicate progressive disruption of cell–matrix adhesion and cytoskeletal integrity. For dermal fibroblasts, attachment to the extracellular matrix provides essential survival signals through integrins and focal adhesion-associated pathways; loss of these interactions can trigger detachment-induced apoptosis, or anoikis [37]. Consequently, the morphological deterioration and increased apoptosis observed at later time points are likely to be mutually reinforcing rather than independent phenomena.

The flow-cytometric findings further support a temporal shift from adaptation to cell death. Cells remained predominantly viable between Day 0 and Day 6, whereas early and late apoptotic fractions increased after Day 9. Annexin V detects the externalization of phosphatidylserine, an early feature of apoptosis, while propidium iodide identifies cells that have lost plasma membrane integrity [38]. The concurrent elevation of Annexin V-positive/PI-negative and Annexin V-positive/PI-positive populations is therefore compatible with the progressive passage of cells from early apoptosis to late apoptosis or secondary necrosis. Nevertheless, Annexin V/PI profiles alone do not define the molecular pathway responsible for death, and phosphatidylserine exposure is not entirely specific to apoptosis under all experimental conditions. Additional measurements of mitochondrial membrane potential, cytochrome *c* release, caspase-3/7 activation, PARP cleavage, BCL-2 family proteins, lipid peroxidation, and intracellular reactive oxygen species would help determine whether sustained 15 °C exposure primarily activates mitochondrial apoptosis, death receptor-associated pathways, oxidative injury, or a combination of these mechanisms.

The transcriptomic data provide molecular evidence that the response to 15 °C is progressive and strongly dependent on exposure duration. Principal component analysis showed that PC1 and PC2 accounted for 71.58% and 19.56% of the total variance, respectively, indicating that the first two components together captured more than 91% of the overall expression variation. Day 0 samples were clearly separated from all hypothermia-exposed samples, while Day 3 and Day 6 were positioned closer to one another and Day 9, Day 12, and Day 15 became increasingly separated along the principal trajectory. PCA is an unsupervised method and therefore does not by itself identify the biological factors responsible for sample separation[39]. Nevertheless, when considered together with the coordinated deterioration in morphology, metabolic activity, and survival, the ordered distribution strongly suggests that treatment duration was a dominant source of transcriptional variation. The proximity of Day 3 and Day 6 also corresponds to their relatively preserved phenotype, whereas the greater divergence of the later samples is consistent with a transition toward injury and cell death. This concordance between independent phenotypic and molecular measurements strengthens the interpretation that prolonged 15 °C exposure causes cumulative biological remodeling rather than random variation among culture samples.

Time-series analysis of differentially expressed genes (DEGs) provided quantitative evidence for dissecting the molecular response dynamics of canine dermal fibroblasts under sustained 15 °C hypothermic stress. Using Day 0 as the baseline, the total number of DEGs exhibited a near-monotonic increasing trend with prolonged exposure: 3,098 on Day 3, rising to 4,092 on Day 6, reaching 4,667 on Day 9, increasing further to 5,086 on Day 12, and peaking at 5,579 on Day 15. This continuously expanding DEG repertoire indicates that hypothermia-induced transcriptomic perturbation was not a single acute event, but rather a progressively accumulating and intensifying process over time.

From a stage-specific perspective, downregulated genes outnumbered upregulated genes on Days 3 and 6, suggesting that early hypothermic stress imposed broad and systematic suppression of fundamental cellular functions, including cell adhesion, cytoskeletal remodeling, energy metabolism, and PI3K-Akt survival signaling. By Day 9, the number of upregulated genes accelerated (from 1,966 to 2,328, an 18.4% increase), whereas the growth of downregulated genes markedly slowed (from 2,126 to 2,339, a 10.0% increase), with the two categories approaching parity for the first time (2,328 vs. 2,339). This shift in transcriptomic composition closely paralleled phenotypic observations. Day 9 coincided precisely with the onset of pronounced cellular shrinkage, cytoplasmic vacuolization, and increased refractivity, as well as the inflection point where CCK-8 metabolic activity shifted from increase to decline and apoptotic fractions began to rise significantly. Thus, Day 9 marks the molecular boundary at which fibroblasts transition from early adaptation to late cumulative injury.

In the late phase (Days 9–15), transcriptomic characteristics underwent more profound changes. Between Days 9 and 12, upregulated genes continued to increase rapidly (+400), while downregulated genes nearly stagnated (+19), resulting in upregulated genes exceeding downregulated genes for the first time (2,728 vs. 2,358). By Day 15, upregulated genes (2,995) significantly surpassed downregulated genes (2,584). The emergence of this “upregulation-dominant” pattern suggests that under severe late-stage hypothermic stress, the intensity of cellular stress responses had exceeded the degree of basal metabolic suppression, with massive induction of late-stage-specific injury-associated genes—such as apoptosis executioners, inflammatory mediators, and innate immune signaling molecules—driving the full establishment of a large-scale stress/damage-response transcriptional program.

These DEG dynamics were corroborated by STEM clustering and GSEA results. Early DEGs likely primarily reflected cold sensing, metabolic suppression, and immediate protective responses; whereas the substantial influx of new DEGs after Day 9 corresponded to progressive activation of innate immune pathways such as autophagy, Toll-like receptor, and RIG-I-like receptor signaling, alongside progressive inhibition of core energy metabolic pathways including glycolysis and the pentose phosphate pathway. This transcriptomic pattern transition—from “suppression-dominated downregulation of basal functions” through “balanced up- and downregulation” to “stress-activation-dominated damage responses”—supports the three-stage response model proposed in this study: early adaptation (Days 0–6), transitional inflection (around Day 9), and late decompensation (Days 12–15).

These findings also have implications for the biology and preservation of primary canine dermal fibroblasts. Fibroblasts are central regulators of extracellular-matrix synthesis, wound contraction, paracrine signaling, and tissue repair[40,41]. Maintenance of cell survival alone is therefore insufficient for preservation applications; retained function after rewarming is equally important. Cells that remain Annexin V-negative after hypothermic exposure may still exhibit growth arrest, senescence, genomic instability, altered collagen production, or impaired migration. Future studies should consequently evaluate clonogenic recovery, population-doubling time, migration, collagen and fibronectin production, myofibroblast differentiation, and cellular senescence after return to physiological temperature. Comparisons with standard hypothermic preservation temperatures, such as 4 °C and 8 °C, and with temperatures closer to routine culture conditions would establish whether 15 °C offers a favorable balance between metabolic suppression and cold injury. Preservation media, cell density, oxygen availability, pH, buffering capacity, and medium renewal should also be optimized because temperature changes alter gas solubility, acid–base balance, nutrient utilization, and waste accumulation. In bicarbonate-buffered media in particular, the relationship between temperature and CO₂ conditions may substantially influence extracellular pH and confound the direct effects of hypothermia.

Several limitations should be acknowledged. First, the study appears to have examined cells from a limited biological background; primary fibroblasts can vary according to donor age, sex, breed, biopsy location, passage number, and health status. Validation in cells from multiple dogs is essential for distinguishing general canine responses from donor-specific effects. Second, CCK-8 measures metabolic reduction rather than proliferation directly, and Annexin V/PI analysis should be supplemented with mechanistic markers of cell death. Third, bulk RNA sequencing averages signals across heterogeneous cell populations and cannot resolve subpopulation-specific stress responses. Fourth, differential expression does not demonstrate that a particular gene or pathway is causally responsible for survival or injury. Candidate regulators identified by trend analysis and functional enrichment should be confirmed using quantitative PCR and protein-level assays, followed by loss- or gain-of-function experiments. Finally, post-hypothermic recovery was not evaluated. Such experiments are indispensable for defining the practical preservation window because cells may appear viable during cold exposure but fail to re-enter the cell cycle or restore normal function after rewarming.

## 4. Materials and Methods

### 4.1. Animals, ethics statement, and skin tissue collection

Skin tissue was harvested from one healthy 18-month-old male local-bred beagle dog housed at the Animal Experiment Center of Naval Medical University. Before tissue collection, the health status of the donor was evaluated by routine physical examination, blood routine and serum biochemical-index laboratory testing to rule out skin disease, systemic infection and underlying organic lesions. Full-thickness skin specimens were collected from the inner thigh skin under aseptic conditions during routine sterile biopsy procedures.

All procedures involving animals were performed in accordance with the relevant institutional and national guidelines for the care and use of animals. The experimental protocol was reviewed and approved by the Institutional Animal Care and Use Committee of Naval Medical University. Skin specimens were immediately transferred into sterile phosphate-buffered saline (PBS) supplemented with 100 U/mL penicillin and 100 µg/mL streptomycin and transported to the laboratory at room temperature within 30 minutes after collection.

### 4.2. Isolation and culture of primary canine dermal fibroblasts

Primary canine dermal fibroblasts were isolated from skin tissues using an explant outgrowth method. Briefly, the samples were washed three times with sterile PBS containing 100 U/mL penicillin and 100 µg/mL streptomycin. Subcutaneous adipose tissue, hair follicles, and other visibly non-dermal tissues were removed using sterile scissors and forceps. The remaining dermal tissue was cut into approximately 1–2 mm³ pieces.

Then, the tissue fragments were placed onto flasks and incubated for 2 hours to allow attachment before the careful addition of complete growth medium. Cells were maintained in high glucose Dulbecco’s modified Eagle’s medium (Thermo Fisher Scientific) supplemented with 10% fetal bovine serum (FBS) (Thermo Fisher Scientific), and 1% penicillin–streptomycin. Cultures were incubated at 37 °C in a humidified atmosphere containing 5% CO₂. The medium was replaced every 2–3 days. After reaching approximately 80–90% confluence, cells were detached using 0.25% trypsin– EDTA and subcultured at a ratio of 1:3. Primary fibroblasts at passages 3 were used for all experiments.

### 4.3. Experimental design and prolonged low-temperature treatment

Third-passage canine dermal fibroblasts were seeded into corresponding culture vessels according to the requirements of each subsequent experiment. Cells were incubated at 37 °C for 24 h to allow stable adherence before further treatment. Then, the cultures assigned to low-temperature treatment were transferred to an incubator maintained at 15 °C for up to 15 days. Samples were collected at Day 0, Day 3, Day 6, Day 9, Day 12, and Day 15. Day 0 samples were collected immediately before transfer to 15 °C and served as the baseline control. During low-temperature exposure, cells were maintained in complete medium under 5% CO₂ concentration. The culture medium was refreshed every 3 days. For each time point, 3 independent replicates were prepared.

### 4.4. Morphological examination

Cell morphology was examined on Days 0, 3, 6, 9, 12, and 15 using an inverted bright-field microscope (Nikon). Before imaging, culture vessels were inspected without disrupting either adherent or detached cells. Representative fields were photographed at 200 magnification using identical acquisition settings for all groups. Morphological evaluation focused on cell shape, spreading, membrane appearance, cytoplasmic vacuolization, refractivity, fragmentation, and detachment from the culture surface.

### 4.5. CCK-8 assay of cellular metabolic activity and viability

Cellular metabolic activity was evaluated using a Cell Counting Kit-8 (CCK-8, CK04, Dojindo, Japan) according to the manufacturer’s instructions. Canine dermal fibroblasts were seeded into 96-well plates at a density of 2 10^3^ cells per well in 100 µL of complete medium. After initial attachment, the cells were exposed to 15 °C and examined at the designated time points.

At each time point, 10 µL of CCK-8 reagent was added to each well containing 100 µL of culture medium. The plates were incubated at 37 °C for 1h under conditions consistent across all groups. Absorbance at 450 nm was measured using a SpectraMax 190 microplate reader (Molecular Devices, USA).

### 4.6. Annexin V-FITC/PI apoptosis assay

Cell apoptosis was quantified using an Annexin V-FITC/propidium iodide (PI) apoptosis detection kit (Yeasen Biotechnology, Shanghai, China). At each time point, culture supernatants containing detached cells were collected first. The remaining adherent cells were washed with cold PBS and detached using trypsin without EDTA. Adherent and floating cells were combined, centrifuged at 500×g for 5 min, and washed twice with cold PBS. Approximately 1 × 10 cells were resuspended in 100 µL of 1× binding buffer. The cell suspension was incubated with 5 µL Annexin V-FITC and 10 µL PI for 15 min at room temperature in the dark. Samples were analyzed within 1 h using a CytoFLEX flow cytometer (Beckman Coulter, USA). At least 10,000 single-cell events were acquired for each sample.

Unstained, single Annexin V-FITC-stained, and single PI-stained control groups were used to establish fluorescence compensation and quadrant boundaries. Cell debris was excluded using forward scatter and side scatter parameters, while cell doublets were excluded via FSC-A versus FSC-H gating. Cells were categorized into four groups according to their staining profiles: viable cells (Annexin V⁻/PI⁻), early apoptotic cells (Annexin V⁺/PI⁻), late apoptotic or secondary necrotic cells (Annexin V⁺/PI⁺), and necrotic/dead cells (Annexin V⁻/PI⁺). Flow-cytometric data were processed using CytExpert (Beckman Coulter, USA). The same gating strategy was applied to all samples.

### 4.7. RNA extraction and quality assessment

Total RNA was extracted from cells harvested at Day 0, Day 3, Day 6, Day 9, Day 12 and Day 15 using TRIzol reagent (Thermo Fisher Scientific, USA) following the manufacturer’s instructions. Only adherent viable cells were collected for RNA isolation at later time points. Contaminating genomic DNA was digested and eliminated by DNase I treatment.

RNA concentration and purity were determined with a Qubit fluorometer (Invitrogen, USA). RNA integrity was verified on an Agilent 2100 Bioanalyzer system (Agilent Technologies, USA). Qualified RNA samples meeting the following criteria were subjected to library construction: RNA integrity number (RIN) ≥ 7.0, A260/A280 ratio ranging from 1.8 to 2.1, and adequate total RNA mass.

### 4.8. Library preparation and RNA sequencing

Sequencing libraries were constructed using 2 µg of high-quality total RNA from each sample. Polyadenylated mRNA was enriched via oligo(dT) magnetic bead purification to remove ribosomal RNA and other non-coding RNAs. The purified mRNA was randomly fragmented into short fragments, and first-strand cDNA was synthesized using reverse transcriptase with fragmented mRNA as the template. Second-strand cDNA was subsequently generated, followed by a series of standard procedures including end repair, 3′-end A-tailing, adapter ligation, and fragment size selection. Purified cDNA fragments were finally amplified by PCR to complete the construction of transcriptome sequencing libraries.

Library concentration and fragment size distribution were accurately quantified using a Qubit fluorometer (Invitrogen, USA) and an Agilent 2100 Bioanalyzer (Agilent Technologies, USA). Qualified libraries with uniform fragment size and sufficient concentration were subjected to high-throughput sequencing on the Illumina NovaSeq 6000 platform (Illumina, San Diego, CA, USA) with a paired-end 150 bp (PE150) sequencing strategy. Each sample obtained approximately 20 million raw sequencing reads to ensure adequate sequencing depth for subsequent differential gene expression and time-series transcriptome analysis.

### 4.9. RNA-seq data processing and expression quantification

Raw sequencing reads were first subjected to quality evaluation using FastQC (v0.11.9). Adapter contaminations, low-quality reads, and ambiguous bases were systematically trimmed and filtered via fastp (v0.23.2). Post-filter quality verification was performed to guarantee the validity of clean reads for subsequent genomic alignment. Clean reads were mapped to the canine reference genome assembly CanFam3.1 (Canis lupus familiaris) using HISAT2 (v2.2.1). Corresponding gene annotation files were obtained from the Ensembl database (Release 109). Gene-level read counts were quantified using featureCounts (v2.0.3).

To reduce technical noise, genes with extremely low expression were pre-filtered prior to downstream analysis; only genes with a read count ≥ 10 in at least one sample were retained. For subsequent bioinformatic analysis, raw read counts were used for differential gene expression analysis, while fragments per kilobase of transcript per million mapped reads (FPKM) normalized values were applied for gene expression visualization and cross-sample comparison across different time-point samples.

### 4.10. Principal component analysis (PCA)

PCA was performed to comprehensively evaluate the global similarities and discrepancies of transcriptome profiles among samples from six culture time points. Raw gene count matrices were normalized and transformed using the variance-stabilizing transformation (VST) implemented in the DESeq2 package to eliminate the mean–variance dependence of sequencing data. PCA analysis was conducted via the prcomp function in R software. The first two principal components (PC1 and PC2) were visualized using the ggplot2 package, and individual samples were colored according to their culture duration to intuitively display time-dependent transcriptomic clustering patterns. The proportion of total transcriptional variance explained by each principal component was calculated based on corresponding eigenvalues to quantify the degree of sample separation.

### 4.11. Differential gene-expression analysis

Differentially expressed genes (DEGs) were identified using the DESeq2 package (v1.38.3) in R (v4.3.1). Pairwise differential expression analysis was performed by comparing each low-temperature treatment time point against the Day 0 baseline, including five independent contrast groups: Day 3 vs Day 0, Day 6 vs Day 0, Day 9 vs Day 0, Day 12 vs Day 0, and Day 15 vs Day 0. Multiple testing correction was performed using the Benjamini–Hochberg false discovery rate (FDR) method to adjust raw P values.

Genes with an adjusted P value < 0.05 and |log₂ fold change (FC)| ≥ 1 were defined as statistically significant DEGs. Genes with log₂FC > 1 were categorized as upregulated, while those with log₂FC < −1 were categorized as downregulated.

### 4.12. Venn and UpSet intersection analyses

The overlap and time-point specificity of DEGs across the five pairwise comparisons were examined using Venn diagrams and UpSet analysis. Only genes satisfying the same differential-expression threshold in each comparison were included. Venn diagrams were generated using VennDiagram, and UpSet plots were produced using UpSetR package in R.

### 4.13. Time-series expression trend analysis

To systematically classify the dynamic temporal expression patterns of stress-responsive genes during prolonged low-temperature culture, the aggregated DEGs from all five pairwise comparisons were subjected to Short Time-series Expression Miner (STEM) analysis[42]. Prior to trend analysis, FPKM expression values were log₂-transformed and Z-score standardized to eliminate dimensional differences and normalize expression fluctuations across time points. According to the standard STEM workflow, the expression level of each gene at Day 3, 6, 9, 12, and 15 was normalized relative to the Day 0 baseline.

STEM parameters were set as follows: a maximum of eight model profiles, a maximum unit change of 2 between adjacent time points, and 1000 permutation tests for statistical evaluation. Individual genes were assigned to specific model profiles based on the Pearson correlation between their actual temporal expression trajectories and predefined theoretical profiles. Permutation-based testing was applied to correct for multiple comparisons, and model profiles with a corrected P < 0.05 were regarded as statistically significant. Profiles with highly similar temporal variation characteristics were further grouped into major expression clusters to summarize representative gene expression patterns and facilitate subsequent functional enrichment interpretation.

### 4.14. Gene Ontology and pathway enrichment analyses

To explore the potential biological functions and signaling pathways involved in long-term low-temperature stress response, Gene Ontology (GO) and Kyoto Encyclopedia of Genes and Genomes (KEGG) enrichment analyses were separately performed for significantly upregulated and downregulated DEGs using the clusterProfiler package (v4.6.2) in R. All canine Ensembl gene IDs were converted to Entrez gene IDs for functional annotation. For genes lacking sufficient canine genomic annotation information, homologous human orthologs were retrieved via the Ensembl BioMart database to ensure comprehensive enrichment analysis coverage.

GO enrichment analysis was conducted across three standard categories: biological process (BP), cellular component (CC), and molecular function (MF). KEGG pathway analysis was performed to identify significantly perturbed signaling cascades associated with cold stress adaptation. All genes that passed pre-filtering and were eligible for differential expression analysis were defined as the background gene set to avoid genome-wide annotation bias. The Benjamini–Hochberg algorithm was used for multiple testing correction, and GO terms and KEGG pathways with adjusted P < 0.05 were regarded as statistically significantly enriched. Redundant and highly similar functional terms were further simplified using the built-in semantic similarity simplify function of clusterProfiler. The top 10 enriched GO terms and KEGG pathways were visualized via bar plots and bubble plots to exhibit the core functional alterations induced by prolonged low-temperature stimulation.

### 4.15. Statistical analysis

Quantitative experimental data from CCK-8 cell viability assays and flow cytometry detection were presented as the mean ± standard deviation (SD) of at least three independent biological replicates. All statistical analyses were performed using GraphPad Prism 9.0 and R software (v4.3.1). P values less than 0.05 were considered statistically significant. Detailed sample sizes, specific statistical methods, and error bar definitions were explicitly stated in the corresponding figure legends.

## 5. Conclusions

In conclusion, continuous culture at 15 °C elicits a dynamic, stage-dependent response in primary canine dermal fibroblasts. A relatively stable early period is followed by a transition beginning around Day 9 and culminating in extensive structural injury, metabolic impairment, apoptosis, and transcriptomic divergence by Day 15. The coexistence of persistent and stage-specific DEGs suggests that prolonged hypothermia engages both a conserved core stress program and temporally distinct adaptive and injury-associated responses. These results establish a useful framework for identifying molecular markers of cold tolerance, defining a safe low-temperature preservation window, and developing interventions that delay the transition from adaptation to irreversible damage. Future work integrating pathway validation, metabolic and mitochondrial assays, multi-donor experiments, and functional recovery after rewarming will be required to translate these findings into robust protocols for the preservation and transportation of primary canine fibroblasts.

## Author Contributions

Conceptualization, J.H., B.Y., and Q.D.; Methodology, Y.W., E.S., A.H., J.H., Y.L. and E.L.; Software, Y.W. and E.S.; Validation, J.H., B.Y. and Q.D.; Formal Analysis, Y.W., E.S., A.H. and J.H.; Investigation, Y.W., E.S., A.H., J.H., Y.L. and E.L.; Resources, B.Y. and Q.D.; Data Curation, E.L. and Y.L.; Writing – Original Draft Preparation, Y.W., E.S., A.H., J.H., and E.L.; Writing – Review & Editing, B.Y., and Q.D.; Visualization, Y.W., E.S., A.H., and E.L.; Supervision, B.Y.; Project Administration, B.Y., and Q.D.; Funding Acquisition, B.Y.All authors have read and agreed to the published version of the manuscript.

## Institutional Review Board Statement

Not applicable.

## Informed Consent Statement

Not applicable.

## Data Availability Statement

The original contributions presented in this study are included in the article. Further inquiries can be directed to the corresponding authors.

